# Novel multicellular prokaryote discovered next to an underground stream

**DOI:** 10.1101/2020.12.25.424384

**Authors:** Kouhei Mizuno, Mais Maree, Toshihiko Nagamura, Akihiro Koga, Satoru Hirayama, Soichi Furukawa, Kenji Tanaka, Kazuya Morikawa

**Author notes:** Deceased (Soichi Furukawa).

## Abstract

The emergence of multicellularity is a key event in the evolution of life and is an attractive challenge among researchers, including those investigating the artificial design of cellular behavior^1^. Multicellular organisms are widely distributed on Earth, and retracing the specific conditions conducive for the initial transition from unicellularity to multicellularity is difficult. However, by examining organisms that inhabit unique (e.g., isolated) environmental niches, we may be able to get a glimpse into primitive multicellularity in the context of a given environment. Here we report the discovery of a new bacterium that displayed multicellular-like characteristics and behavior. The bacterium, which was isolated adjacent to an underground stream in a limestone cave, is to be named *Jeongeupia sacculi* sp. nov. HS-3. On a solid surface, HS-3 self-organizes its filamentous cells to form an appearance similar to the nematic phase of a liquid crystal^2^. Mature colonies produce and accommodate clusters of coccobacillus progeny, and release them upon contact with water. HS-3 demonstrated novel, spatiotemporally regulated multicellularity that can resolve the so-called ‘competition-dispersal trade-off’ problem^3^. This study illustrates a hypothetical missing link on the emergence of multicellularity.

## Introduction

The emergence of multicellularity is a mystery of life on Earth, and while it appears to have occurred both repeatedly and independently in different lineages, the generative process is unknown^4,5,6^. The origin of multicellularity has attracted the attention of scientists in a wide range of fields, including bioengineering of artificial cellular behavior^1^ and cancer progression^7^. Multicellularity in prokaryotes is not directly linked to the origin of animals, but it provides us with fundamental models for considering the evolutionary steps of how and why unicellular organisms made the leap to multicellularity. Types of extant prokaryotic multicellularity are summarized in Supplementary Fig. 1. The ‘clonal multicellularity’^5^ is defined as being self-organized through serial cell division without separation of daughter cells. It is commonly seen in animals and some prokaryotes such as actinomycetes and cyanobacteria (Supplementary Fig. 1). These bacteria produce reproducible and patterned structures of clonally developed cells which increase the fitness of the whole population, irrespective of whether or not it may benefit some individuals^8^. In the evolutionary step to clonal multicellularity, certain ancestral modes of ‘group life’^9^ have been hypothesized to represent the intermediate status, but no one has ever observed this missing link (Supplementary Fig. 1).

The cave microbiome was previously considered to be simply a subset of surface microbes^10^, but it has recently been shown that the overlap of operational taxonomic units (OTUs) with microbes from the surface is only 11–16%^11, 12^. Furthermore, the microbiomes of different cave environments differ, as has been revealed in caves composed of limestone^11, 13, 14^, lava^12^, ice^15^, quartz^16^, or sulfur^17^. Namely, each cave has a distinctive microbiome that has adapted to that environment.

Here we report a new cave bacterium that can develop a multicellular architecture from a simple, but ordered, cell cluster, which is reminiscent of ‘group life’. The bacterium was isolated from the surface of a cave wall above an underground river in a limestone cave system that is submerged intermittently. The most remarkable aspect of this bacterium’s life-cycle is the existence of well-regulated dimorphism between the first growth stage (filamentous cells that show liquid crystal-like self-organization on a solid surface) and the second stage (coccobacillus cells that can disperse in flowing water). The second stage strongly suggests the involvement of recurrent water flows in the establishment of the multicellularity in this species. The discovery of this new species supports and further augments the hypothesis that recurrent environmental dynamism is a driving force to the emergence of multicellularity^9^.

## Results

### *Jeongeupia sacculi* sp. nov. HS-3, a new bacterial species isolated from a karst cave

We have collected oligotrophic bacteria from a variety of environments and studied their physiology^18, 19, 20^. In 2008, strain HS-3 was isolated from water dripping on a limestone cave wall in the Hirao karst plateau^21^ in northern Kyushu Island, Japan (Supplementary Fig. 2). The sampling site, which was located slightly above the water surface of an underground river, was intermittently submerged due to an increase in runoff after rainfall (Fig. 1a, Supplementary Fig. 2). After plating on agar, colonies appeared transparent and iridescent (Fig. 1b, Supplementary Fig. 3). HS-3 is a gram-negative obligate aerobe with a single polar flagellum (Fig. 1f, inset). Genetically, it belongs to a group in the family *Neisseriaceae* that comprises mainly environmental bacteria (Supplementary Fig. 4). It has 3.4 Mbp genome and a plasmid (2.0 kbp) (Supplementary Fig. 5). Based on phenotypic comparisons with other closely related species (Supplementary Table 1–3), HS-3 is considered to be a novel species. The optimal growth temperature (24°C) and pH (8.0–9.0) are consistent with conditions in the cave. Another species in the same group that is adapted to life in oligotrophic environments is *Deefgea rivuli*, which has been isolated from tufa, a form of limestone^22^ (Supplementary Fig. 4). However, no reports of distinctive colony morphology or cell differentiation have been reported in this group to date.

**Figure 1.**
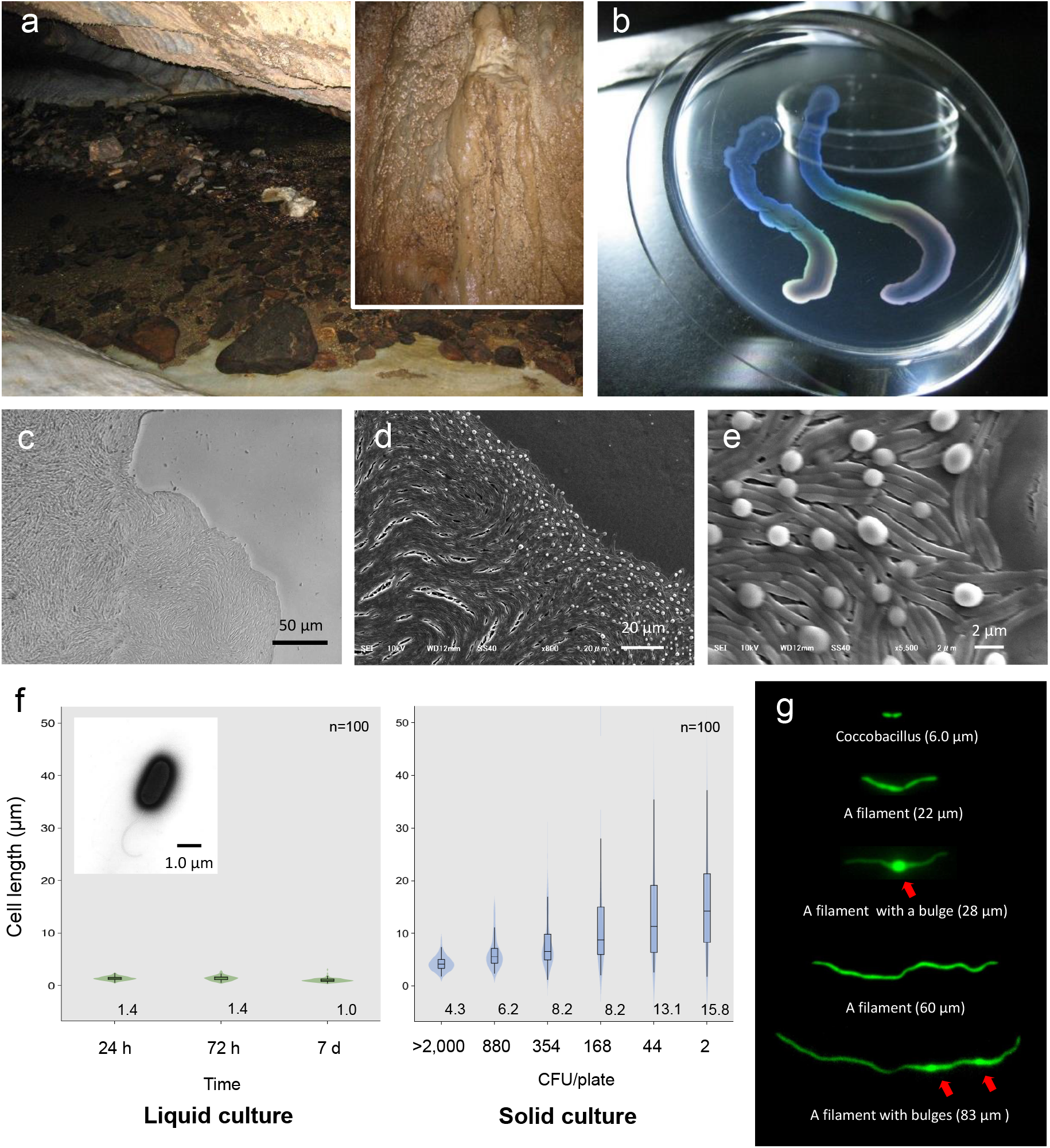
Liquid-crystal appearance and micro-structure of a colony of *Jeongeupia sacculi* sp. nov. HS-3, discovered in a limestone cave. **a**, River in a cave showing the sampling site on the cave wall (inset). **b**, Transparent and iridescent appearance of colonies. **c–e**, The edge of a colony growing on agar. Differential interference contrast (DIC) (**c**) and scanning electron microscopic (SEM) (**d, e**) images. Bulges of cells on the colony surface (**e**). **f**, Cell length in liquid and solid cultures. The liquid cultures were monitored at different times. The solid cultures were inoculated at different colony densities (CFU/plate) and cultured for 2 days. The value given for each sample is the mean obtained from 100 cells. Inset: Transmission electron microscopic (TEM) image of a liquid-cultured cell. **g**, Typical cell morphology from a colony cultured on agar and observed by GFP fluorescence. Arrows: Bulges of cells.

### HS-3 forms colonies with a liquid crystal texture

Bacterial cells capable of growth on agar medium typically proliferate in a disordered state, yielding colonies with an opaque texture. HS-3 formed transparent colonies with an iridescent hue (Fig. 1b). Specifically, the colonies assumed an anisotropic optical structure (Supplementary Fig. 3a), suggesting that the cells were orientated in a manner similar to liquid crystal molecules. As shown in Fig. 1c–e, the cells in the colonies formed vortex-shaped structures composed of closely arrayed filamentous cells with bulges on the surface. Transforming these cells with the green fluorescent protein (GFP) gene revealed that these structures were composed of cells rather than some extracellular matrix or artifacts derived from electron microscopy (Fig. 1g). On a solid medium, the length of the cells was correlated with the colony density, and lower densities were associated with a wider range of cell length (Fig. 1f). These findings suggest that cell elongation in this strain is a regulated process that occurs in response to environmental cues. Cell elongation was not observed in liquid cultures.

A time course analysis of the growing colonies showed that the cells can start to proliferate as coccobacilli for the first 10 h (Supplementary Movie 1). Then, as cell elongation was initiated from the edge of the colony (Supplementary Movie 2), the cells gradually assumed topological characteristics similar to the two-dimensional nematic pattern described in liquid-crystal theory^23^, with comet-like (+1/2) and trefoil-like (−1/2) topological defects being visible (Supplementary Fig. 6). Generally, the microcolonies initially spread as a single layer. As the colony expands, bulges are produced especially, but not specifically, around the rapidly growing colony edge (Fig. 1e). We consider that the morphological change occurs to sustain the ordered state by relieving internal physical pressure. The disturbed single layer at the center then serves as the focal point for the expansion of additional layers (Fig. 2ab, Supplementary Fig. 7). The bulges are also generated around the colony center (Supplementary Fig. 8). As the multilayered colony continues to grow, the expanding edge swallows the adjacent region and the internal filamentous cells buckle, generating domains with a vortex structure (Fig. 2e, Supplementary Movie 3). Importantly, the filamentous cells appear to sustain the well-aligned structure, suggesting that there is tight adhesion among the filamentous cells (Fig. 2e). An anisotropic pattern that emerged on the bottom layer of the colony was sustained throughout the period of colony growth (Fig. 2cd). Furthermore, the optical characteristics of the multi-layer colonies indicate that the ordered structures are sustained (Supplementary Fig. 3a).

**Figure 2.**
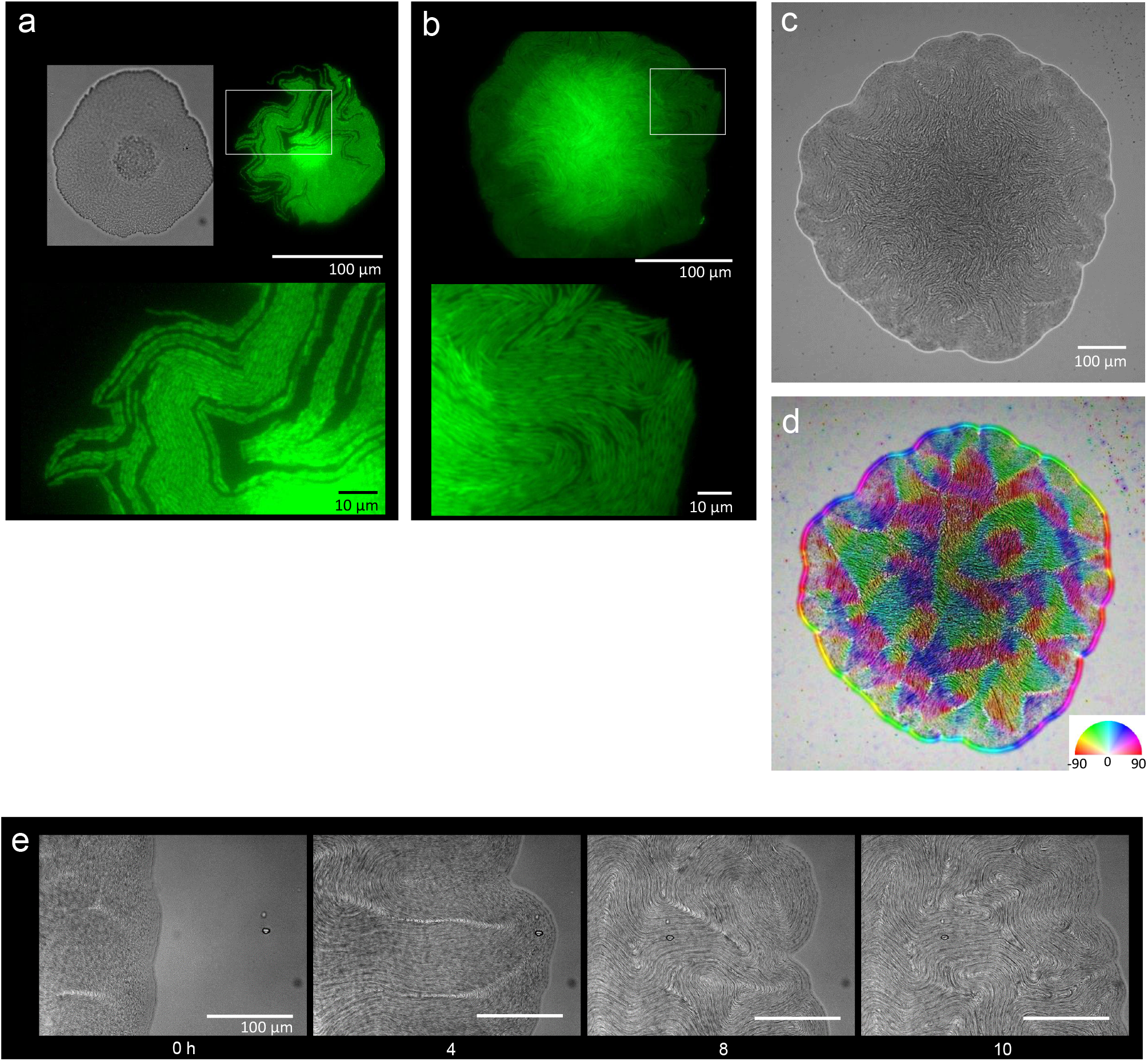
Colony morphogenesis and liquid crystal-like ordering. **a**, A young colony showing a single layer of cells. DIC (top left) and GFP (top right) images and an enlarged image (bottom) of the area indicated by the white box. **b**, A microcolony with a single layer of cells at the edge. Enlarged image of the area indicated by the white box (bottom). **c**, Anisotropic pattern of the bottom cell layer of a single colony. **d**, The color mapping to show the orientation of cells. The anisotropic pattern is a typical characteristics in a nematic phase^2^. **e**, Time-lapse images of liquid crystal-like pattern formation on the edge of a growing colony. Time (h) after chase is indicated. The original movie is available as **Supplementary Movie 3**.

### Internal proliferation to germ-like phase

Colonies stopped growing at 2 days after inoculation and no further changes were observed for the following 2-3 days. However, internal proliferation then started and the colony began to swell three-dimensionally (Supplementary Fig. 9a, Supplementary Movie 4). Internally proliferating cells formed optically distinct regions within a transparent colony, suggesting the loss of the ordered-layer structure. Nile red, a lipid-staining fluorescent dye, was used to selectively stain the membranes of the elongating cells in a colony^24^, and the internal coccobacillus cells were recognized as non-stained areas in three-dimensional cross-sectional images (Fig. 3a), as well as in micrographs obtained by conventional fluorescence microscopy (Fig. 3b, Supplementary Fig. 9b). At the 5th day after inoculation, the internal cells were crowded-out of the colony (Fig. 3b right panels). Interestingly, the occurrence of this event in a single colony triggered a chain reaction of this phenomenon in adjacent colonies (Fig. 3c), indicating that the crowding out of coccobacilli is under some form of control.

**Figure 3.**
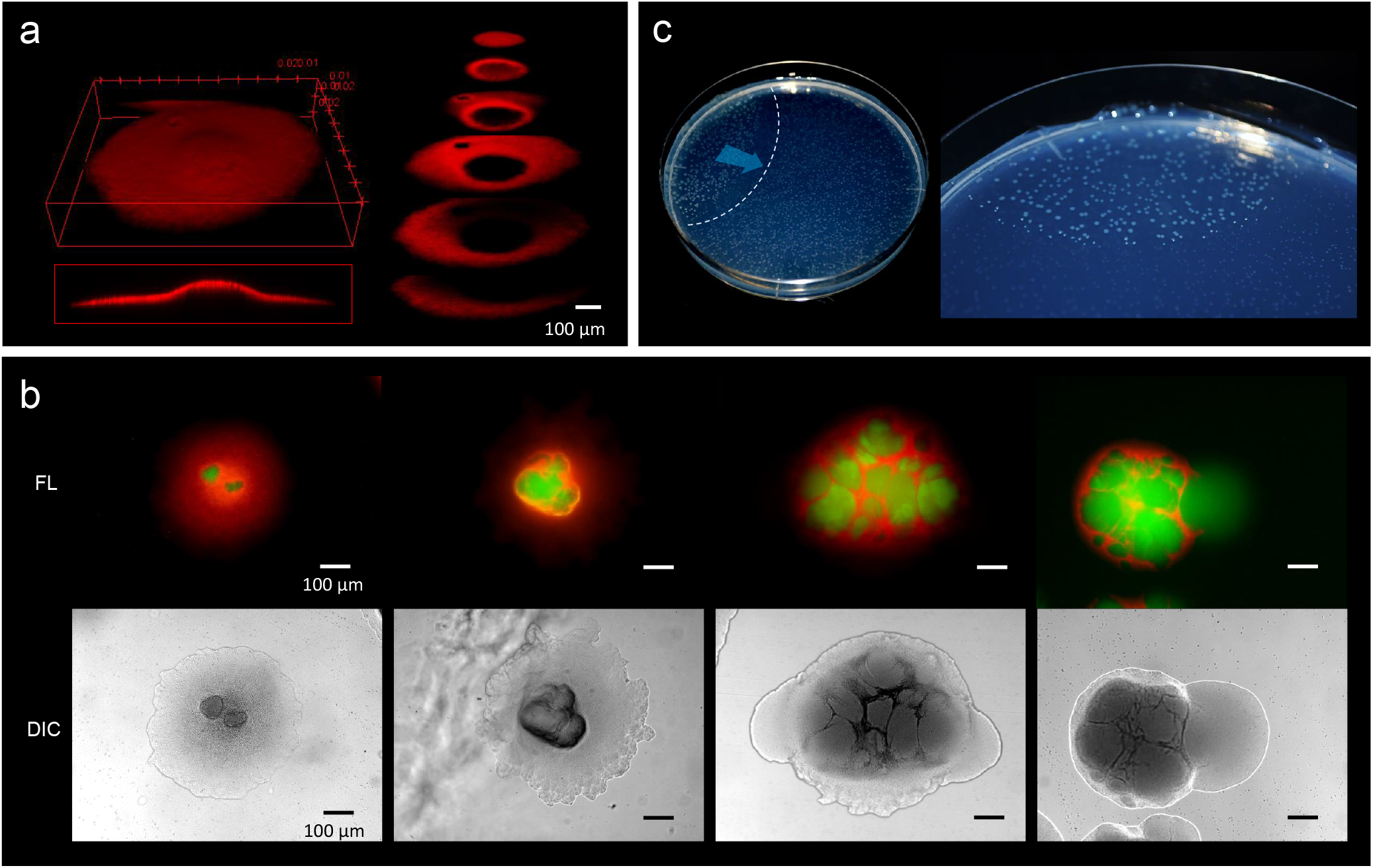
Internally growing cells and crowding out. **a**, Three-dimensional image of a 4-day-old colony reconstructed from 100 Z-axis slices with a total height of 168 μm. The whole Nile-red image (left) and its cross sections (right). **b**, Five-day-old, mature colonies showing internal growth at the center of the colony. Two right panels show the ‘crowding out’. **C**, Colony-crowding influences adjacent colonies by inducing a ‘chain reaction’. The direction of the chain reaction and its front line are indicated by the blue arrow and the dotted line, respectively. More details of a crowding colony are shown in **Supplementary Movie 4**.

The sampling site of HS-3 was a moist cave wall above an underground river (Fig. 1a). Since this site is also intermittently submerged by water due to increased runoff due to rainfall, we examined the response of colonies on an agar plate to being submerged by water. Remarkably, the coccobacilli were released into the water column, leaving the architecture produced by the filamentous cells behind (Fig. 4a, Supplementary Movie 5). Even after the complete release of the coccobacillus from the center of the colony, the bottom cell layer of the aligned filamentous cells adhered tightly to each other (Fig. 4a, Supplementary Fig. 10). These observations suggest that the filamentous cells act to support the ‘spawning’ of the embedded coccobacilli clusters. We also examined whether the spawned daughter cells were able to reproduce the original colony morphology. The results showed that the spawned coccobacillus cells could indeed reproduce the original filamentous colony morphology (Supplementary Fig. 11a). In addition, when filamentous cells obtained from a mature colony were inoculated on agar (Supplementary Fig. 11b, Supplementary Movie 6), they became elongated and then divided to generate short rods that each formed a micro-colony. Thus, the filamentous cells were not irreversibly destined to be the equivalent of ‘somatic’ cells. These results suggest that the different morphological changes observed in the life-cycle of HS-3 are reversible (Fig. 4b).

**Figure 4.**
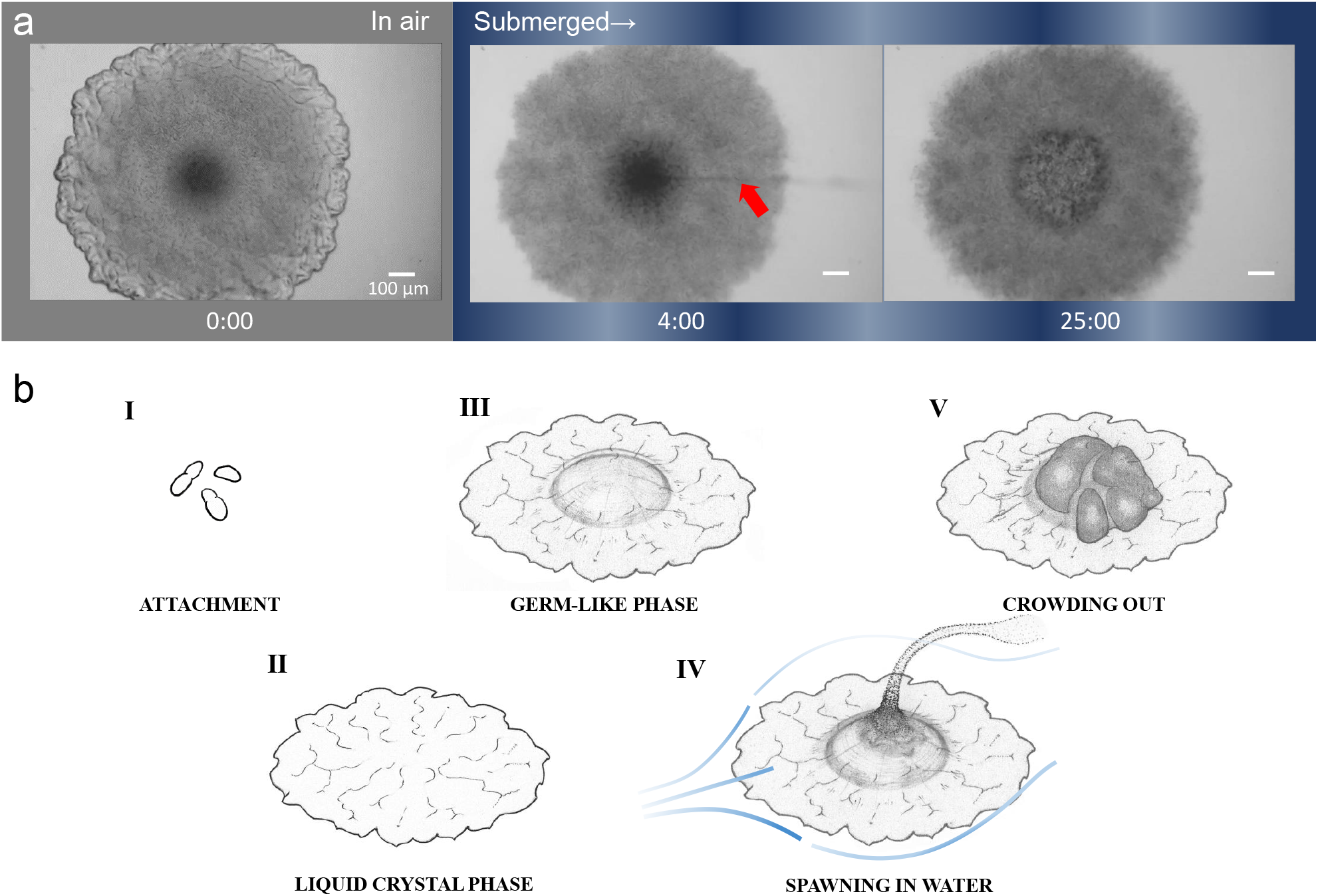
Waterborne cells and multicellular life-cycle of HS-3. **a**, Time-lapse images of a mature colony in air (left panel) and after being submerged in water (middle and right panels). The red arrow indicates the waterborne coccobacillus cells released from the colony. The colonies were grown for 5 days on agar before the time-lapse analysis. The time (min) after chase is shown. Bars = 100 μm. The original movie is available as **Supplementary Movie 5**. **b**, Illustration of the multicellular life-cycle of HS-3. (**I**) Coccobacillus cells attached to a new solid surface. (**II**) The cells proliferate to some extent, generating a liquid-crystal texture. (**III**) The center area becomes raised due to clustering of internal cells, leading to the germ-like phase. (**IV**) A waterflow releases coccobacilli. (**V**) In the absence of a waterflow, outward growth of the internal coccobacillus cells results in ‘crowding out’.

## Discussion

The physiological significance of multicellularity has been attributed to avoiding predation^25^, cooperative defense^4^, labor division^4, 5, 26^, and creation of a stable internal environment^5^ (Supplementary Fig. 1). While little is known about why and how some single-celled organisms became multicellular, an underlying driving mechanism for this transition is considered to be recurrent environmental dynamics^9^. Indeed, the importance of recurrent environmental dynamics is supported by both experiments using model organisms^3, 27^ and theoretical predictions^9^. Strain HS-3 utilizes both episodic flooding of the river or the continuous supply of water dripping on the cave wall to efficiently disperse their progeny.

It is considered that hypothetical ancestors employing a ‘group life’ must have made a leap in structural order^28^ towards multicellularity (See the “Missing Link: ‘Group life’“ in Supplementary Fig. 1). Some pioneering experiments using model organisms^27, 29^ have proposed that a ‘dichotomy’ in the ‘group life’ mode is a prerequisite for the emergence of multicellularity. The concept of ‘dichotomy’ is defined as having two distinct, but co-existing, cellular characteristics in a group of cells^29^. Such characteristics can be observed in extant single-celled bacteria, which can produce differentiated subpopulations using a variety of genetic (e.g., phase variation) or non-genetic (e.g., bistability) mechanisms^30, 31^. By producing these differentiated subpopulations, they can achieve division of labor or hedge their bets in the event of unpredictable future conditions^32^. Although these mechanisms include dichotomic characteristics, the organisms have not yet acquired distinctive multicellular structures; Note that “Differentiation & subpopulation” is not included in ‘Clonal multicellularity’ (Supplementary Fig. 1). Namely, dichotomy alone is not sufficient for the establishment of structurally ordered multicellularity.

HS-3 employed cellular dichotomy comprising a structural filamentous framework and the emergence of coccobacillus progeny to achieve a multicellular architecture. Another pivotal characteristic of HS-3 is that the emergence of germ cells was regulated to be set aside until maturation of the filamentous framework, rather than being a spontaneous event. This study hypothesizes that sequential ordering (e.g., in HS-3, the delayed generation of germ-like cells following the establishment of nematic liquid-crystal-type layers), in addition to dichotomy, is a prerequisite for developing multicellularity.

## Acknowledgements

We thank the following people for their support with this study: Dr. Masahiro Nakano for discussion on aspects related to physics. Ms. Aya Ohta for isolating the bacterium, Ms. Satoko Nakanomori and Mr. Sakae Fukase for phenotype characterization, and Mr. Hiroyuki Tanaka for SEM analysis.

## Materials and methods

### Strains and culture media

Strain HS-3 was isolated in 2008 from percolating water on the surface of a limestone cave wall in ‘Seryukutsu’ (Blue Dragon) cave in the Hirao karst plateau (33°46’00.5”N 130°55’02.7”E) in northern Fukuoka Prefecture, Japan (Supplementary Fig. 2a–c). The sampling site on the limestone wall was situated close to the surface of an underground river. Water dripping from the wall was collected using a sterile plastic tube and the pH and temperature were measured on site. Samples were then stored at 4°C for several days until analysis. The water sample was diluted with phosphate-buffered saline (PBS, pH 7.0) to inoculate R2A (0.5 g yeast extract, 0.5 peptone, 0.5 casamino acids, 0.5 glucose, 0.3 K_2_HPO_4_, 0.05 MgSO_4_ per liter of deionized water) agar plates containing Nile red (Sigma-Aldrich, Missouri, USA) at 0.5 mg per liter. The HS-3 strain was originally isolated as a Nile red-positive strain, indicating that it was a lipid producer. The isolated HS-3 strain was pre-cultured in R2A liquid medium in a test tube overnight at 24°C in a reciprocal shaker at 100 rpm. Then, the culture was diluted with PBS and inoculated on a plate containing R2A and 1.5% agar for taxonomic and microscopic analysis. The close relatives, *Jeongeupia naejangsanensis* KCTC 22633^T^ and *J. chitinilytica* KCTC 23701^T^ were purchased from the Korean Collection for Type Cultures, Taejon, Korea. Two closely related bacteria, *Andreprevotia chitinilytica* NBRC106431^T^ and *Silvimonas terrae* NBRC100961^T^, were purchased from the National Bio-Resource Center, Kisarazu, Japan. Those strains were cultured in R2A medium at 24 or 30°C on agar, or in a reciprocal shaker at 100 rpm for liquid cultures.

### Taxonomic identification of strain HS-3

The Gram reaction was performed using a Gram Staining Kit (Sigma). Culture media and conditions for taxonomic comparisons were essentially the same as those used for the reference strains, *J. naejangsanensis* KCTC 22633^T 33^ and *J. chitinilytica* KCTC 23701^T 34^. Growth at various temperatures (4, 10, 16, 18, 20, 22, 24, 26, 28, 30, 32, 34, and 40°C) and pH (3.0–11.0 at intervals of 0.5) was performed in nutrient broth (Becton Dickinson, New Jersey, USA). The pH was adjusted by using sodium acetate/acetate, phosphate, and Na_2_CO_3_ buffers. Growth at different NaCl concentrations (0, 0.5, 1.0, 2.0, 3.0, 4.0, and 5.0% w/v) was also performed in tryptic soy broth prepared according to the formula used for the Difco medium, except that NaCl was excluded. Growth under anaerobic conditions was performed using an anaerobic chamber with a gas exchange kit (Anaeropack; Mitsubishi Gas Chemical, Tokyo, Japan). Catalase and hydrolysis of casein, gelatin, starch, and Tween 20, 40, 60, and 80 were determined as described by Cowan & Steel^35^. Additionally, assimilation of Tween 20, 40, 60, and 80 as sole carbon sources was tested in M9 medium (12.8 g Na_2_HPO_4_·7H_2_O, 3 g KH_2_PO_4_, 0.5 g NaCl, and 1 g NH_4_Cl per liter of deionized water containing 2 mM MgSO_4_ and 0.1 mM CaCl_2_). Susceptibility to antibiotics was tested by placing antibiotic-impregnated discs on agar plates seeded with the test strain. The antibiotics tested were kanamycin, gentamycin, erythromycin, fosfomycin, trimethoprim, novobiocin, streptomycin, chloramphenicol, ampicillin, and tetracycline. Utilization of various substrates and biochemical properties were tested by using API 20NE, API 50CH, API ZYM kits (bioMerieux, Marcy l’Etoile, France). Major types of quinones was determined at Techno Suruga, Shizuoka, Japan, using reverse–phase high-performance liquid chromatography (HPLC)^36^. Cellular fatty acid composition was also determined at Techno Suruga using the MIDI/Hewlett Packard Microbial Identification System^37^. The strain was deposited at the National Bioresource Center, Kisarazu, Japan as *<u>Jeongeupia</u>* sp. HS-3 NBRC 108274^T^ (=KCTC 23748^T^).

Genomic DNA was extracted using a Qiagen kit (Qiagen, Hilden, Germany), and the 16S rRNA gene was amplified using the primer set of 27f and 1525r^38^. The PCR product was sequenced (Applied Biosystems 3730xl DNA Analyzer, Applied Biosystems, USA), and the DNA G + C content was measured by Techno Suruga Inc. using HPLC^39^. A phylogenetic tree was constructed using the neighbor-joining method^40^ implemented in the Clustal W software package^41^ using sequences of closely related bacteria in GenBank that were selected by BLAST^42^.

Genomic DNA for genome sequencing was extracted using a NucleoSpin kit (Macherey-Nagel, Düren, Germany). Complete sequencing was performed at the Oral Microbiome Center, Takamatsu, Japan, using a combination of long-read sequencing performed using a Nanopore system (Oxford Nanopore Technologies, Oxford, UK) and short-read sequencing performed using a DNBSEQ system (MGI Tech, Shenzhen, China). The sequence data were analyzed using Lasergene software (DNASTAR) and annotations were performed by DFAST (https://dfast.nig.ac.jp/). The complete genome sequence and a plasmid sequence have been registered in the DDBJ database (https://www.ddbj.nig.ac.jp/) as AP024094 and AP024095, respectively.

### GFP recombination

Competent cells of strain HS-3 were prepared using general laboratory protocols^43^. Briefly, cells cultured in liquid R2A medium for 48 h at 24°C were inoculated into a fresh R2A medium at a dilution of 20 times. Then, the cells were cultured until the OD_600nm_ reached 0.5. The cells were harvested by centrifugation and washed 2–3 times with sterile water and 10% glycerol, with a gradual increase in the concentration of the suspended cells. Finally, the cells were suspended in 10% glycerol, distributed to tubes, and stored at −80°C. The broad-host-range plasmid, pBBR1MCS2-pAmp-EGFP^44^ (Nova Lifetech, Hong Kong, China), was transformed into competent cells of HS-3 by electroporation under the following conditions. A mixture containing 150 ng of the plasmid and 100 μl of the cell suspension was placed in a cuvette with a 2.0 mm gap and a voltage of 1.8 kV was applied. The suspension was then mixed with 1 ml of R2A medium and incubated for 2 h at 24°C with shaking. The transformants were selected on an R2A agar plate containing kanamycin (25 μg/mL).

### Light and fluorescence microscopy

Cells were cultured in R2A medium either in a test tube containing liquid medium, or on a 1.5% agar or 3.0% gellan gum plate. The liquid-cultured cells were then observed on a glass slide under a microscope (Nikon Eclipse 80i, Nikon Inc., Japan). For time-lapse video microscopy, colonies grown on agar were observed directly on a Petri dish under the microscope using a 10× or 40× objective. Nile-red fluorescent images were obtained for the colonies that were inoculated and grown as described above. The colonies were observed under the microscope with filters for excitation (525–540 nm) and emission (> 565 nm) wavelengths. To obtain three-dimensional images of a colony, a small piece of agar was cut out from an agar plate and placed on a glass slide. The slide was then observed under a confocal laser scanning microscope (Leica Microsystems TCS SP8, Leica Microsystems, Wetzlar, Germany) with filters for excitation (488–552 nm) and emission (560–620 nm) wavelengths and a 10× objective. The images were processed using ImageJ 1.52 software (https://imagej.nih.gov/ij/).

### Electron microscopy

A gel slice of a 48-h-old colony was cut out from a Petri dish and soaked overnight in a 4% paraformaldehyde solution in 100 mM phosphate buffer (pH 7.4), and then soaked for 1 h twice in 20 mM phosphate buffer (pH 7.0). The slice was then dried in a desiccator at room temperature overnight, and then dried under reduced pressure (−20 kPa) overnight. The slice was mounted on an appropriate grid and examined under a scanning electron microscope (JSM-6510LA, JEOL, Tokyo, Japan). Transmission electron microscopy was performed at Hanaichi Ultrastructure Research Institute, Nagoya, Japan. The liquid-cultured cells were mounted on a grid and washed with distilled water at room temperature. Excess liquid was removed and the specimen was negatively stained using 2% uranyl acetate before being observed by TEM (JEM1200EX, JEOL).

