## Supplementary Figures and Tables for "Novel multicellular prokaryote discovered next to an underground stream"

### Supplementary Tables

**Supplementary Table 1 | Characteristics that distinguish strain HS-3 from its close phylogenetic relatives.**

**Supplementary Table 2 | Cellular fatty acid contents (%) of strain HS-3 and its close phylogenetic relatives.**

**Supplementary Table 3 | Antibiotics susceptibilities of strain HS-3 and *J. naejangsanensis*.**

### Supplementary Figures

**Supplementary Figure 1 | Overview of prokaryotic multicellularity and the 'missing link' of the evolutionary step.**

**Supplementary Figure 2 | The isolation site of strain HS-3.**

**Supplementary Figure 3 | Transparent and iridescent colony of HS-3.**

**Supplementary Figure 4 | Phylogenetic tree based on 16S rRNA gene sequence.**

**Supplementary Figure 5 | Genome and plasmid information of HS-3.**

**Supplementary Figure 6 | Emergence of analogous nematic patterns on the colony edge.**

**Supplementary Figure 7 | Initial morphogenesis of a young colony.**

**Supplementary Figure 8 | Internal structures and bulges in a colony.**

**Supplementary Figure 9 | Growth in cargo and spawning tissue.**

**Supplementary Figure 10 | The center area of a colony after contact with water.**

**Supplementary Figure 11 | Reversibility between filament and coccobacillus cells.**

### Supplementary Movies

##### Supplementary Movie 1 | Initial proliferation on agar.

The total of 24 h at  intervals 2–3 min.

##### Supplementary Movie 2 | The edge of a colony growing on solid medium.

The total of 15 h at  intervals 20 min.

##### Supplementary Movie 3 | The edge of a colony further growing on solid medium.

The total of 30 h at  intervals 10 min.

##### Supplementary Movie 4 | Crowd-out of coccobacillus cells from a mature colony.

The total of 33 h at  intervals 10 min.

##### Supplementary Movie 5 | Waterborne coccobacillus cells from a colony.

The movie was edited at 16-folds speed. The actual recording time was 7 min.

##### Supplementary Movie 6 | Reversible proliferation of a mature filament cell on agar.

The total of 5.5 h at  intervals 2–3 min.

### MULTICELLULAR PROKARYOTES

**Clonal multicellularity**[5] describes a mode of multicellularity that arises from a hypothetical ancestral '**group life**'[9] consisting of cell clumps formed through serial cell division without the separation of daughter cells. On the other hand, other multicellular-like behavior, such as differentitation[32] or aggregation[5], may have been alternative options for single-celled prokaryotes. It is currently difficult to determine how and why an ancestral single-celled prokaryotes adopted clonal multicellular characteristics.

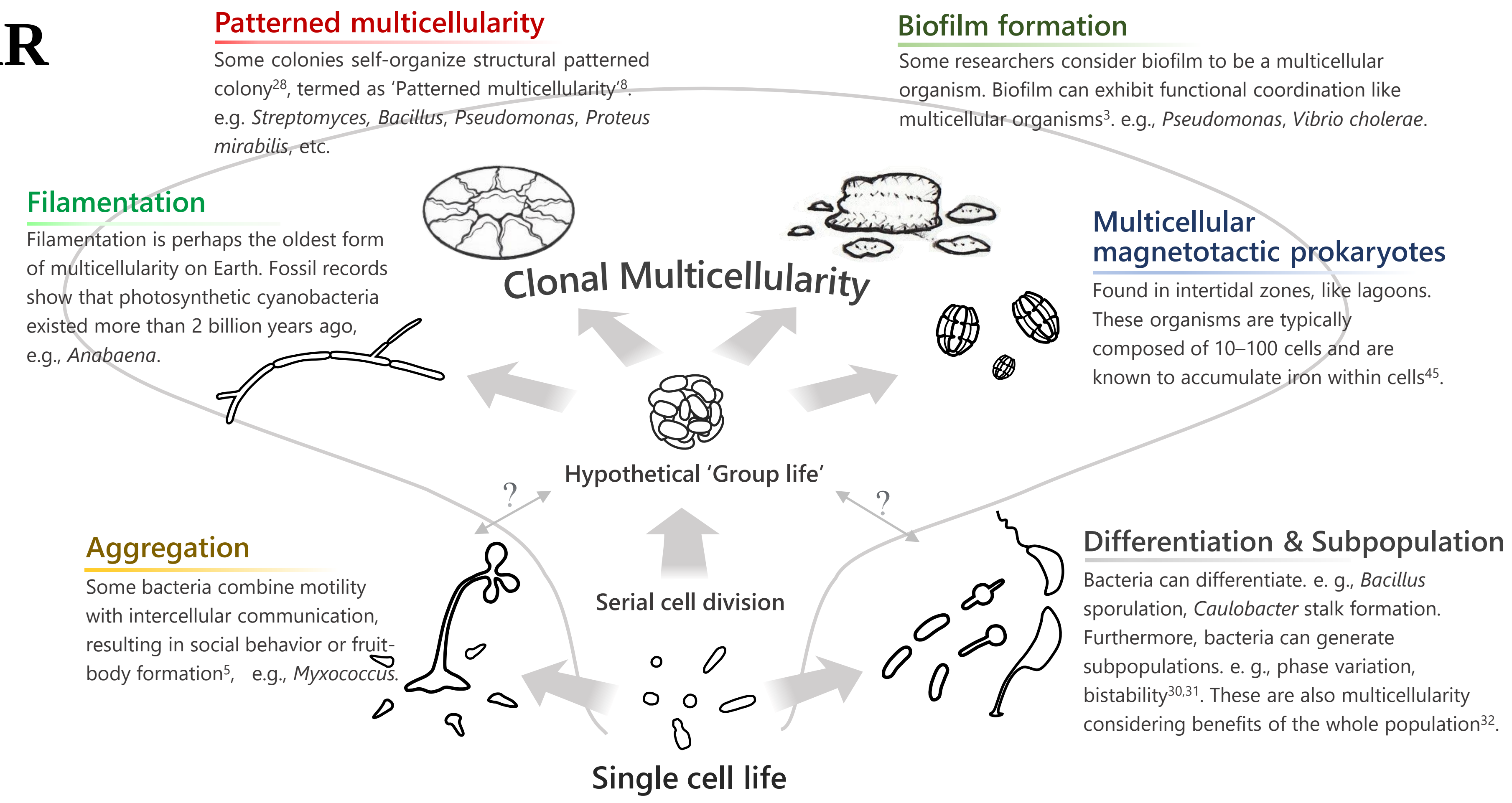

### MISSING LINK: Hypothetical 'Group life'

The 'group life' is a hypothetical intermediate form that represents the transition from unicellularity to multicellularity. However, no one has ever seen such an organism or missing link.

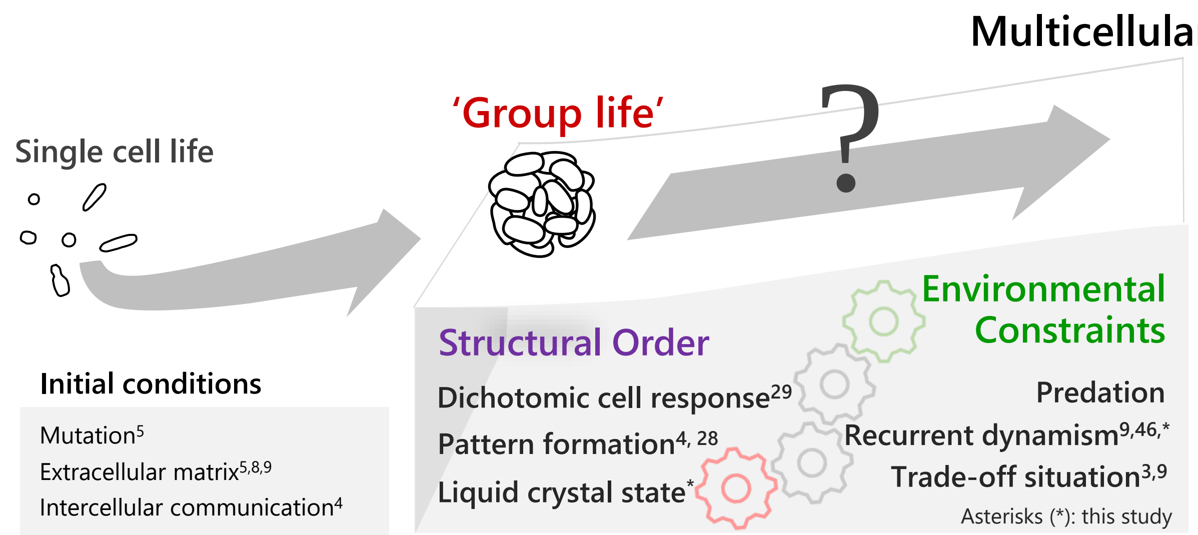

Panel 1: **Phenotypes in current multicellular organisms**

| Term |
|---|
| Spore or cyst formation[8] |
| ECM (extracellular matrix)[8,9] |
| PCD (programmed cell death)[47,48] |
| Intercellular communication[4] |

Panel 2: **Adaptive benefits**

| Term | Description |
|---|---|
| Collective defense against antagonists[4] | e.g., Biofilm against oxidative damage, antibacterial agents, etc. |
| Population survival[4] | Increased chance of differentiation, adaptive DNA exchange or mutation, etc. |
| Cooperative system[5,8] | Behaviors for greater goods. e. g., Altruism like PCD, differentiation for fruiting body or sporulation. |
| Division of labor[4,5,26] | Segregation of somatic/germ function (swimming/reproduction) in volvocine algae. |

Panel 3: **Hypothetical conditions of emergence of multicellularity.**

| Term | Description |
|---|---|
| **Recurrent dynamism**[9,27] | Environmental constraint should reoccur to support the reproducibility of the first group formation across generations. e.g., Feast/famine cycle[9]. Two-phase life cycle in an *in vitro* experiment using *Pseudomonas fluorescence*[27]. |
| **Dichotomic cell response**[8,29,32] | Segregation seen in group life such as soma/germ or reproducing/PCD types. |
| **Trade-off situation**[3,8,9,32] | Ancestral group life should often face such a situation. e. g., 'Competition and dispersal' trade-off situation. |

The emergence of multicellularity has been studied based on phenotypes of extant multicellular organisms (**Panel 1**). Although the adaptive benefits of multicellularity have been proposed for many organisms (**Panel 2**), and are summarized in **Panel 3**, this evolutionary step remains a mystery (**'Group life'**). It is believed that if some form of self-ordering emerges autonomously, it is then selected for in order to overcome a particular environmental constraint. Some conditions are described in **Panel 3**. However, no one has ever observed this process or an intermediate species because it either happened too long ago, or because it could be a one-off event. Resolving this mystery will require a breakthrough that likely involves the discovery of a new model organism.

**Supplementary Figure 1 | Overview of prokaryotic multicellularity and the missing link of the evolutionary step.**

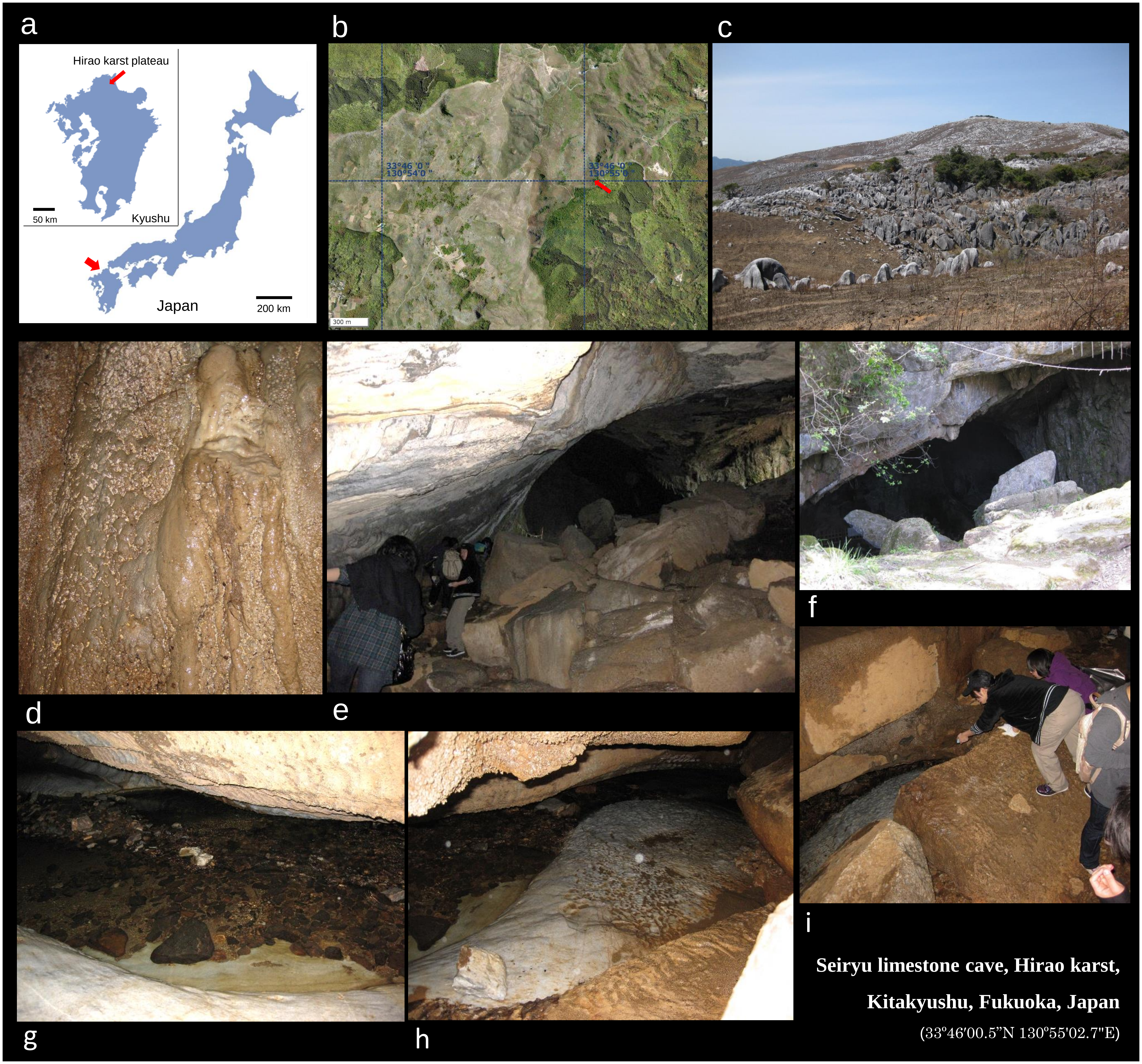

**Supplementary Figure 2 | The isolation site of strain HS-3.**

Images of the site where the bacterium was collected in a limestone cave; **a,** Map of Japan. Arrowhead: Kyushu Island. (Inset) Arrowhead: Hirao karst plateau; **b,** Aerial photograph of the sampling site in the Hirao karst plateau, the whole area of which measures 6.5 km from north to south and 2.5 km from east to west, and has an altitude of between 300 and 700 m. Arrowhead: Seiryu limestone cave. The aerial image was obtained from Geospatial Information Authority of Japan; **c,** Photograph of Hirao karst plateau; **d,** Cave wall (sampling site); **e,** Entrance viewed from inside the cave; **f**, Entrance from outside the cave; **g–i**, River flowing in the cave. The air temperature inside the cave was around 15 ºC. The pH of the water on the wall was 8.5.

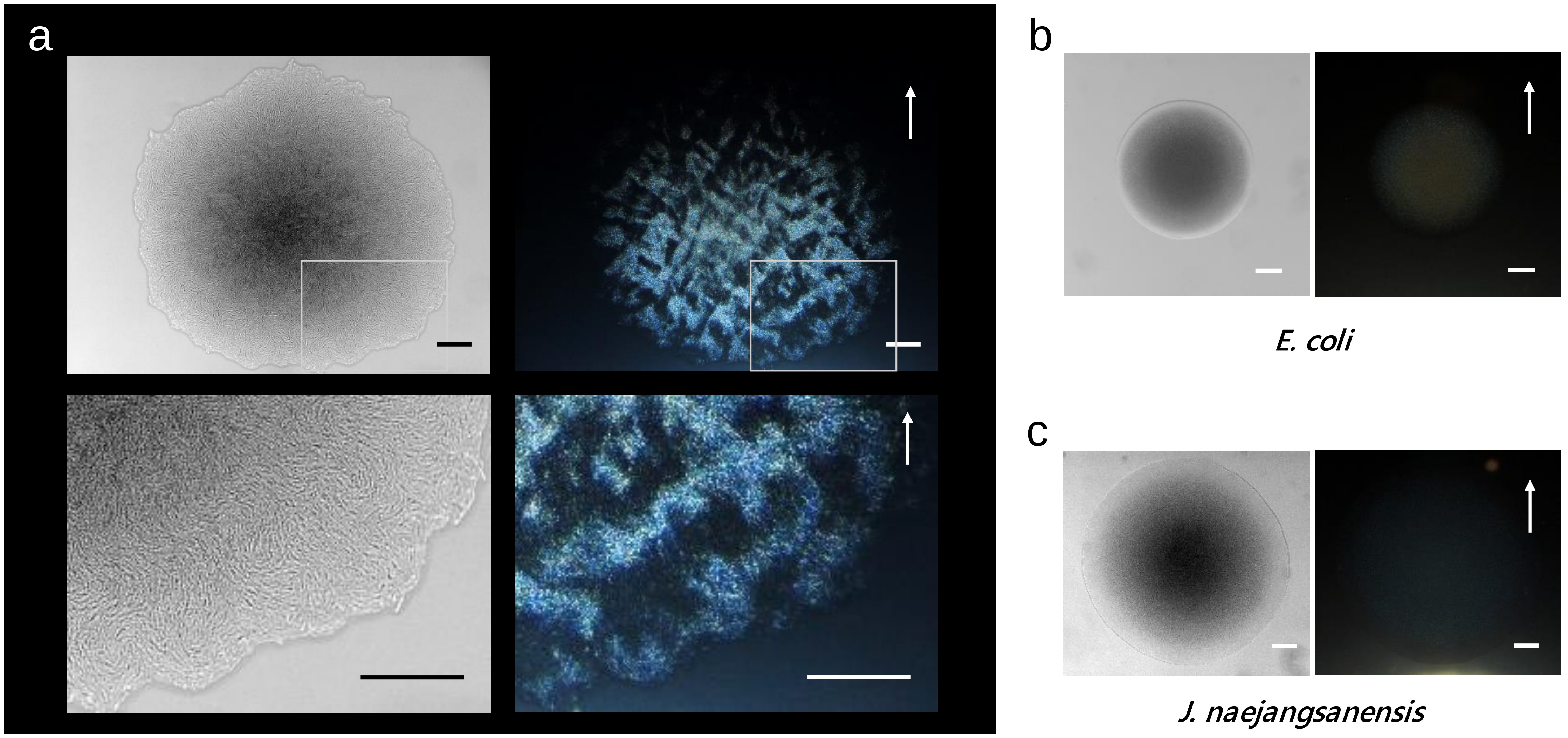

**Supplementary Figure 3 | Transparent and iridescent colony of HS-3.**

Optical microscope images. DIC images (left) and diagonally illuminated images (right). Arrows indicate the direction of light illumination. Bars = 100 µm.

**a**, A colony of HS-3 and a magnified image of the area in the box.
**b**, *E. coli*
**c**, *Jeongeupia naejangsanensis*

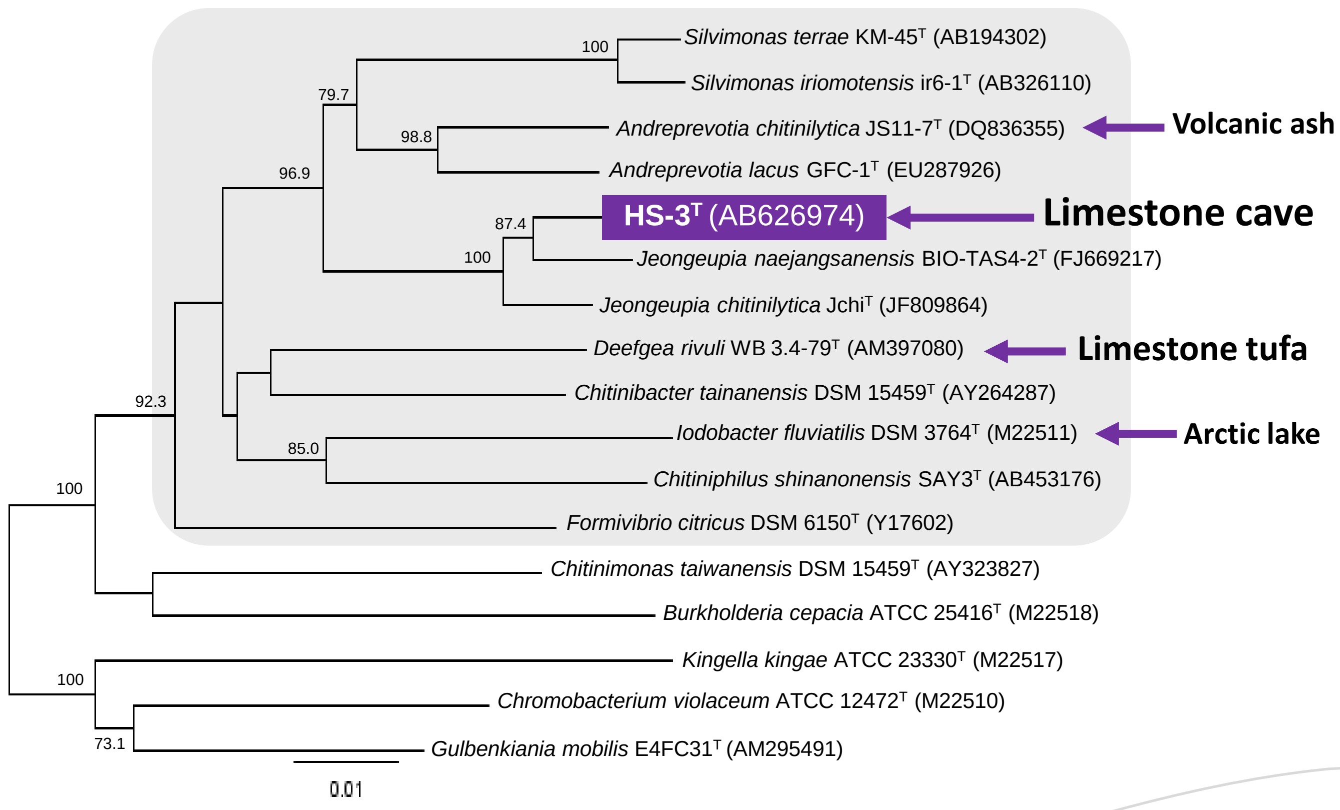

#### Supplementary Figure 4 | Phylogenetic tree based on 16S rRNA gene sequence.

The tree represents a cluster in the *Neisseriaceae* family (shadowed area) and some relatives isolated from similar oligotrophic environments such as tufa, another form of limestone, as indicated by blue arrows. The closest relatives of strain HS-3$^T$ are *Jeongeupia naejangsanensis* KCTC 66233$^T$ (FJ669217) and *Jeongeupia chitinilytica* Jchi$^T$ (NR_109420) with similarity values of 98.2 and 97.8%, respectively.

a

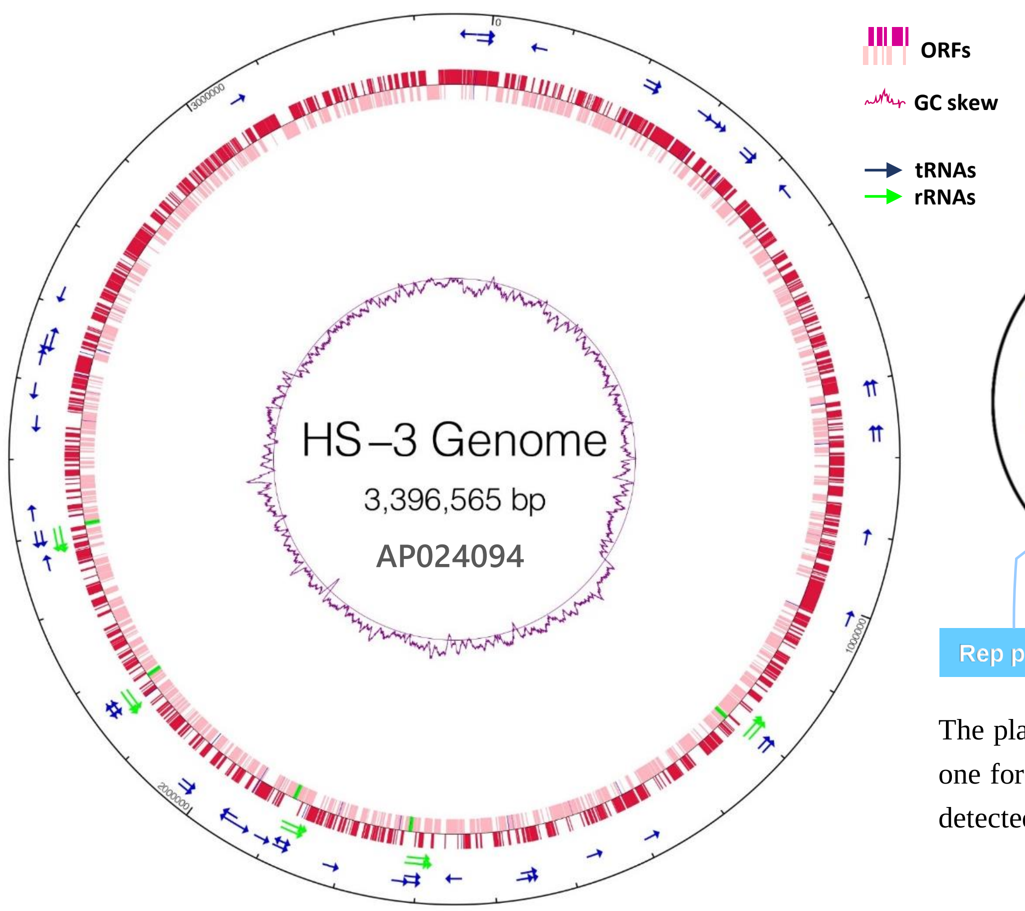

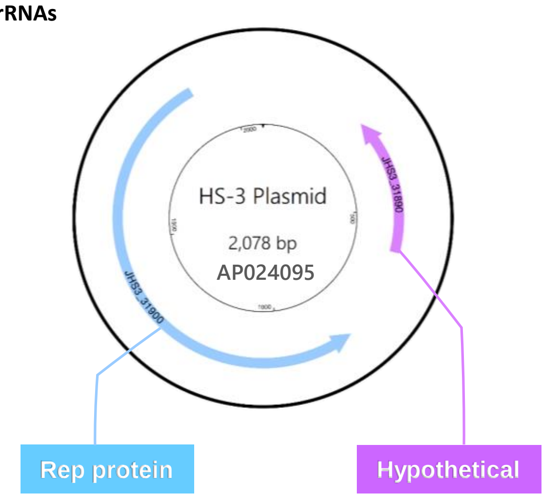

The plasmid seems to be a low copy number one for which it has not been experimentally detected by a commercial extraction kit.

**Panel 1: Summary of the genome of HS-3.**

| Total Sequence Length | |
|---|---|
| Chromosome (bp) | 3,396,565 |
| Plasmid (bp) | 2,078 |
| Number of CDSs | 3,190 |
| GC content (%) | 62.1 |
| Coding Ratio (%) | 89.3 |
| Number of rRNAs | 15 |
| Number of tRNAs | 62 |
| Number of CRISPRs | 1 |

b

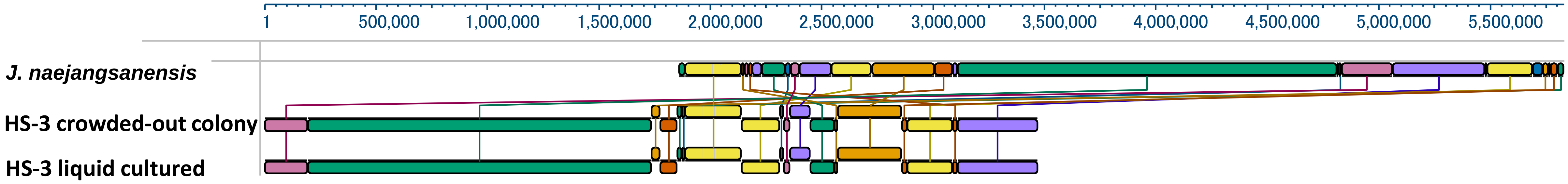

#### Supplementary Figure 5 | Genome and plasmid information of HS-3.

**a,** Diagrams of the genome and the plasmid.

**b,** Pairwise genome comparison among culture conditions and the closest relative.
The analysis was performed using the MAUVE algorithm, which shows large-scale evolutionary events.
The results show that liquid-cultured and crowded-out cells were genetically identical.

#### Supplementary Table 1 | Characteristics that distinguish strain HS-3 from its close phylogenetic relatives.

+, Positive; −, negative; W, weakly positive. All taxa are positive for motility, catalase, acid phosphatase, N-acetyl-β-glucosamidase activity, nitrate reduction, and utilization of α-D-glucose, D-mannose, and N-acetyl-β-glucosamine. All taxa are negative for gram-staining as well as alkaline phosphatase, lipase (C14), valine arylamidase, cyctine arylamidase, trypsin, α-chymotrypsin, α-galactosidase, β-galactosidase, β-glucosidase, α-mannosidase, and α-fucosidase activity. All three strains have Q-8 as their major quinone.

**Reference data**: *J. naejangsanensis* KCTC 22633$^T$ (Yoon *et al*., 2010)[33]; *J. chitinilytica* Jchi KCTC 23701$^T$ (Chen *et al*., 2013)[34].

| Characteristic | *J. sacculi* sp. nov. HS-3$^T$ | *J. naejangsanensis* | *J. chitinilytica* Jchi |
|---|---|---|---|
| **Shape** | Rods or filament | Rods | Rods |
| **Singly or in pairs** | Singly or in pairs | Singly | Singly |
| **Cell size** ( μm, width × length) | 1.0 × 1.0–1.4 in liquid<br>1.5 × 5.0–80 on solid | 0.3–0.7 × 0.7–2.5 | 0.3-0.5 × 1.0–2.0 |
| **Anaerobic growth** | − | + | + |
| **Growth at 40℃** | − | + | − |
| **Glucose fermentation** | − | + | + |
| **Optimal pH** [on solid] | 6.5 [8.0-9.0] | 7.0–8.0 [6.0-9.0] | 7.0–8.0 |
| **Temperature range** (°C) [optimal] | 16–32 [24] | 10–40*[30] | 20–37 [25–30] |
| **Utilization of :** | | | |
| Adonitol | − | + | + |
| Maltose | − | − | + |
| Xylitol | − | + | + |
| Citrate | − | + | + |
| Malate | − | + | + |
| Caprate | − | + | + |
| Cellulose | − | + | − |
| Tween 80 | + | − | + |
| Glycogen | − | + | − |
| **Enzyme activity** (by API ZYM): | | | |
| Esterase Lipase (C8) | w | w | − |
| Leucine arylamidase | w | + | + |
| Naphthol-AS-BI-phosphohyderolase | w | − | − |
| α-glucosidase | − | − | + |
| **DNA G + C content (mol%)** | 62.1 | 63.8 | 66.1 |

*Fifty in the original description (Yoon *et al*., 2009)

#### Supplementary Table 2 | Cellular fatty acid contents (%) of strain HS-3 and its close phylogenetic relatives.

—, ＜0.5% or not detected.
**Reference data**: *J. naejangsanensis* KCTC 22633$^T$ (Yoon *et al*., 2010)[33]; *J. chitinilytica* Jchi KCTC 23701$^T$ (Chen *et al*., 2013)[34].

| Fatty acid | *J. sacculi* sp. nov. HS-3$^T$ | *J. naejangsanensis* | *J. chitinilytica* Jchi |
|---|---|---|---|
| **Straight-chain** | | | |
| $C_{12:0}$ | 3 | — | 1.9 |
| $C_{13:0}$ | 1.2 | — | — |
| $C_{14:0}$ | 5.2 | 8.3 | 7.6 |
| $C_{15:0}$ | 8.7 | 4 | — |
| $C_{16:0}$ | 20.3 | 25.5 | 25.5 |
| $C_{17:0}$ | 1 | — | 0.5 |
| **Unsaturated** | | | |
| $C_{15:1}\omega5c$ | 0.2 | 0.4 | 0.3 |
| $C_{15:1}\omega6c$ | 5.4 | 1.2 | 0.4 |
| $C_{17:1}\omega6c$ | — | 6.3 | — |
| $C_{18:1}\omega7c$ | 3.1 | — | 12.6 |
| **Cyclo** | | | |
| $C_{17:0}$ | 9.7 | — | 8.2 |
| **Hydroxy** | | | |
| $C_{12:0}$ 3-OH | 4.8 | 5.3 | 5.4 |
| **Summed features *** | | | |
| 1 | 0.5 | — | — |
| 3 | 33.7 | 47.1 | 29.0 |

*Summed features represent groups of two or three fatty acids that cannot be separated by GLC with the MIDI system.
Summed feature 1 contains iso-$C_{15:1}$ H and/or $C_{13:0}$ 3-OH. Summed feature 3 contains $C_{16:1}\omega7c$ and/or iso-$C_{15:0}$ 2-OH.

#### Supplementary Table 3 | Antibiotics susceptibilities of strain HS-3 and *J. naejangsanensis*.

Susceptibility to antibiotics was tested by placing antibiotic-impregnated discs on TSA plates that were seeded with suspensions of the test strain. Bars mean no inhibition zone. ND: Not determined.

| Antibiotics | Concn. [ μg/ml ] | *J. sacculi* sp. nov. HS-3$^T$ 24h | *J. sacculi* sp. nov. HS-3$^T$ 48h | *J. naejangsanensis* 24h | *J. naejangsanensis* 48h |
|---|---|---|---|---|---|
| **Kanamycin** | 5,000 | 21.4 | 20.5 | 16.5 | 16.7 |
| | 2,000 | 12.5 | 11.9 | - | - |
| | 1,000 | - | - | - | - |
| | Control | - | - | - | - |
| **Gentamicin** | 5,000 | 36.3 | 36 | 33.55 | 32.6 |
| | 2,000 | 27 | 27.6 | 28 | 27.1 |
| | 1,000 | 26 | 25 | 23.5 | 22.15 |
| | Control | - | - | - | - |
| **Streptomycin** | 5,000 | 26 | 28 | 24 | 26 |
| | 2,000 | 16.7 | 18.4 | 20 | 20.1 |
| | 1,000 | 11.2 | 12 | 16.2 | 16.7 |
| | Control | - | - | - | - |
| **Erythromycin** | 5,000 | 31.1 | 31 | 28.8 | 30 |
| | 2,000 | 23.05 | 22.5 | 24.7 | 24 |
| | 1,000 | 18 | 18 | 20.85 | 21.05 |
| | Control | - | - | - | - |
| **Fosfomycin** | 5,000 | - | - | 17 | 16.2 |
| | 2,000 | - | - | 11.55 | 11 |
| | 1,000 | - | - | - | - |
| | Control | - | - | - | - |
| **Trimethoprim** | 5,000 | 25.5 | 24.5 | 28.2 | 26.7 |
| | 2,000 | 19 | 17.5 | 20 | 19.4 |
| | 1,000 | - | - | 13.4 | 12.5 |
| | Control | - | - | - | - |

| Antibiotics | Concn. [ μg/ml ] | *J. sacculi* sp. nov. HS-3$^T$ 24h | *J. sacculi* sp. nov. HS-3$^T$ 48h | *J. naejangsanensis* 24h | *J. naejangsanensis* 48h |
|---|---|---|---|---|---|
| **Novobiocin** | 5,000 | 37 | 35.2 | 30.55 | 26 |
| | 2,000 | 31.4 | 30 | 25.75 | 22 |
| | 1,000 | 26 | 25 | 22 | 20 |
| | Control | - | - | - | - |
| **Ampicillin** | 4,000 | ND | ND | - | - |
| | 3,000 | ND | ND | - | - |
| | 2,500 | ND | ND | - | - |
| | 2,000 | 22 | 23 | - | - |
| | 1,000 | 14.5 | 15 | - | - |
| | 500 | 13.7 | 13.8 | - | - |
| | 400 | 12.5 | 12.5 | ND | ND |
| | 300 | 11.8 | 11.6 | ND | ND |
| | 200 | - | - | ND | ND |
| | 100 | - | - | - | - |
| | Control | - | - | - | - |
| **Chloramphenicol** | 2,000 | 19 | 18.5 | - | - |
| | 1,000 | 16 | 15 | - | - |
| | 500 | 10.8 | 10 | - | - |
| | 100 | - | - | - | - |
| | Control | - | - | - | - |
| **Tetracycline** | 500 | 16 | 15 | 14.5 | 14 |
| | 400 | 15.5 | 14.8 | 13.8 | 13 |
| | 300 | 13 | 12 | 12 | 11.5 |
| | 200 | 11 | 10.2 | - | - |
| | 100 | - | - | - | - |
| | Control | - | - | - | - |

Diameter of inhibition ring [ mm ]

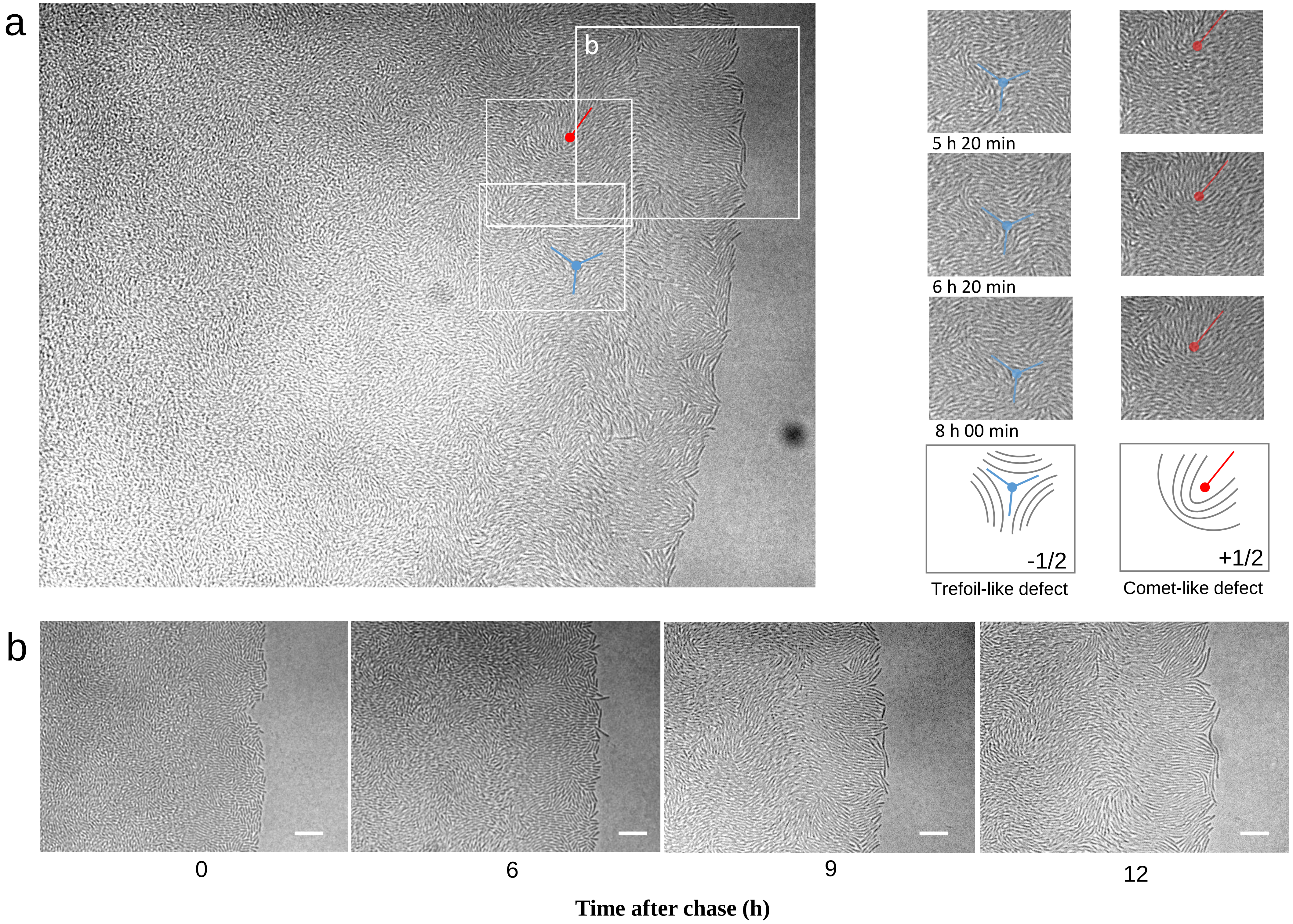

**Supplementary Figure 6 | Emergence of analogous nematic patterns on the colony edge.**

**a**, The emergence of analogous nematic patterns at the colony edge. Two boxed areas are shown as examples on the right. The areas are chronologically aligned and show two types of topological defects characterized by two-dimensional nematic liquid crystals.
**b**, Time course images of the enlarged area indicated by a box on the colony edge.
The original movie is available as **Supplementary Movie 2**.

A key feature underlying the development of the unique cellular architecture is related to the cells adopting the two-dimensional, nematic liquid crystal orientation[2], in which cells are aligned tightly together to form the layered structure. Other bacteria, such as *E. coli* and *B. subtilis*, can produce nematic features, but only for short periods when the cells are allowed to grow two-dimensionally[49,50]. In these species, collisions between the expanding layer of cells results in edge instability and buckling of the two-dimensional nematic order. Conversely, HS-3 filamentous cells have the remarkable ability to sustain an ordered structure.

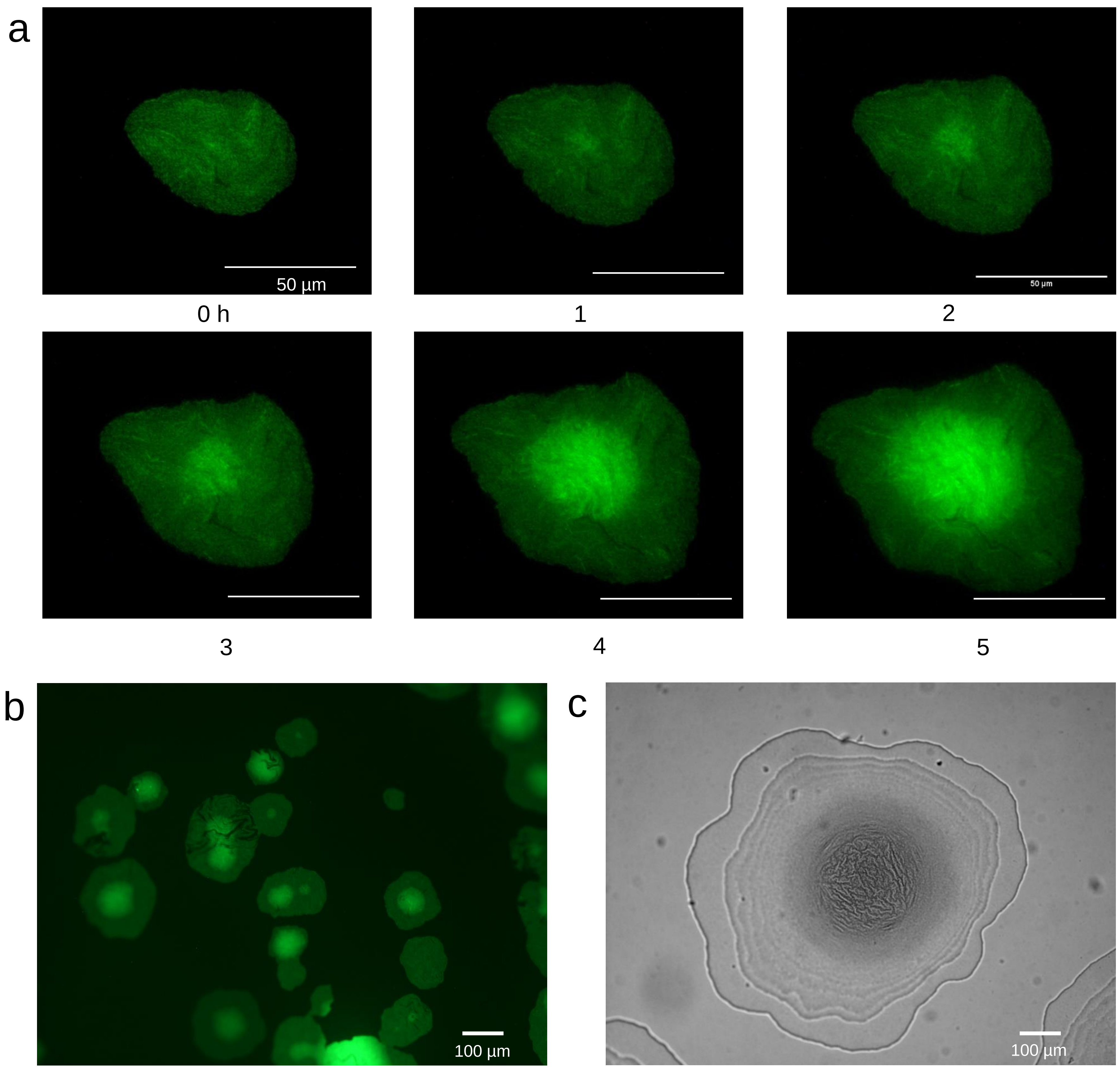

**Supplementary Figure 7 | Initial morphogenesis of a young colony.**

**a**, Time course GFP images of a monolayer colony with  buckling at the center. Time (h) after chase is shown.
**b**, Micro-colonies at the initial stage of the growth.
**c**, A young colony with a ladder structure. The center area shows folds due to bucking.

It appears that HS-3 has the ability to limit the breakage of the monolayer structure to the center of the colony, probably by employing the force directed towards the center and the physical characteristics of the topological defects[2]; it is known that cells tend to gather towards topological defects[50]. Even in the multi-layer state, HS-3 was capable of forming an optically-ordered structure consisting of bundles (Fig. 2d), which we believe may act as rigid framework for the emerging coccobacilli progeny cells.

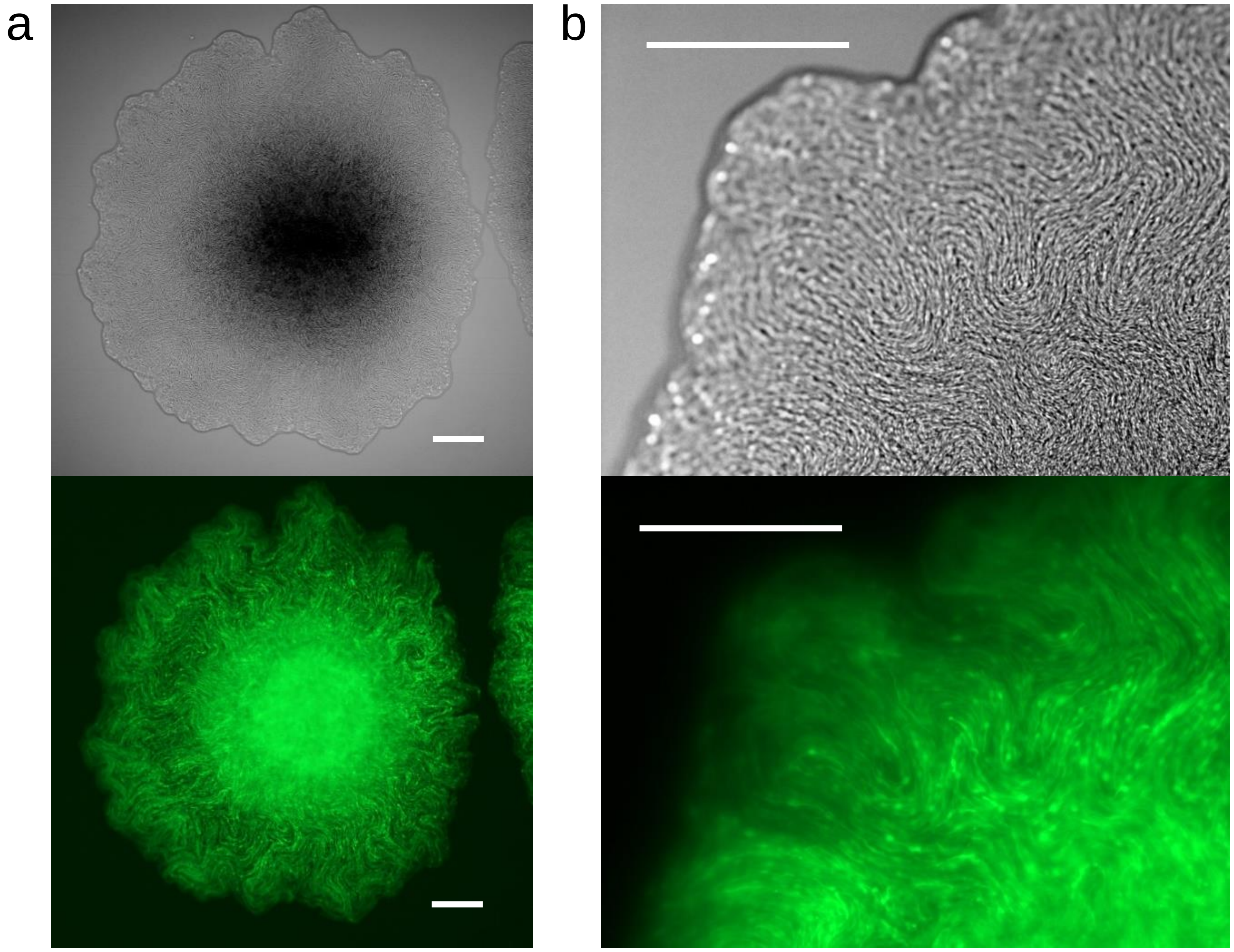

**Supplementary Figure 8 | Internal structures and bulges in a colony.**

DIC and GFP images of a 48 h-old colony. Bars = 100 µm.
**a, b,** The surface showed tightly aligned patterns in the DIC image, while numerous bulges were observed beneath the surface in GFP mode.

HS-3 filamentous cells have the remarkable ability to sustain an ordered structure; this ability must employ a mechanism that can buffer, or relax, the local tension generated by collisions of the elongating and proliferating cells. While we have not clarified the exact mechanism, the formation of the distinct bulges is one candidate; these bulges are generated around the edge of the rapidly growing colony, which would likely be experiencing increased friction on the agar surface (compared to a slow-growing colony at high colony densities). Notably, the bulges are also generated around the colony center (**b**).

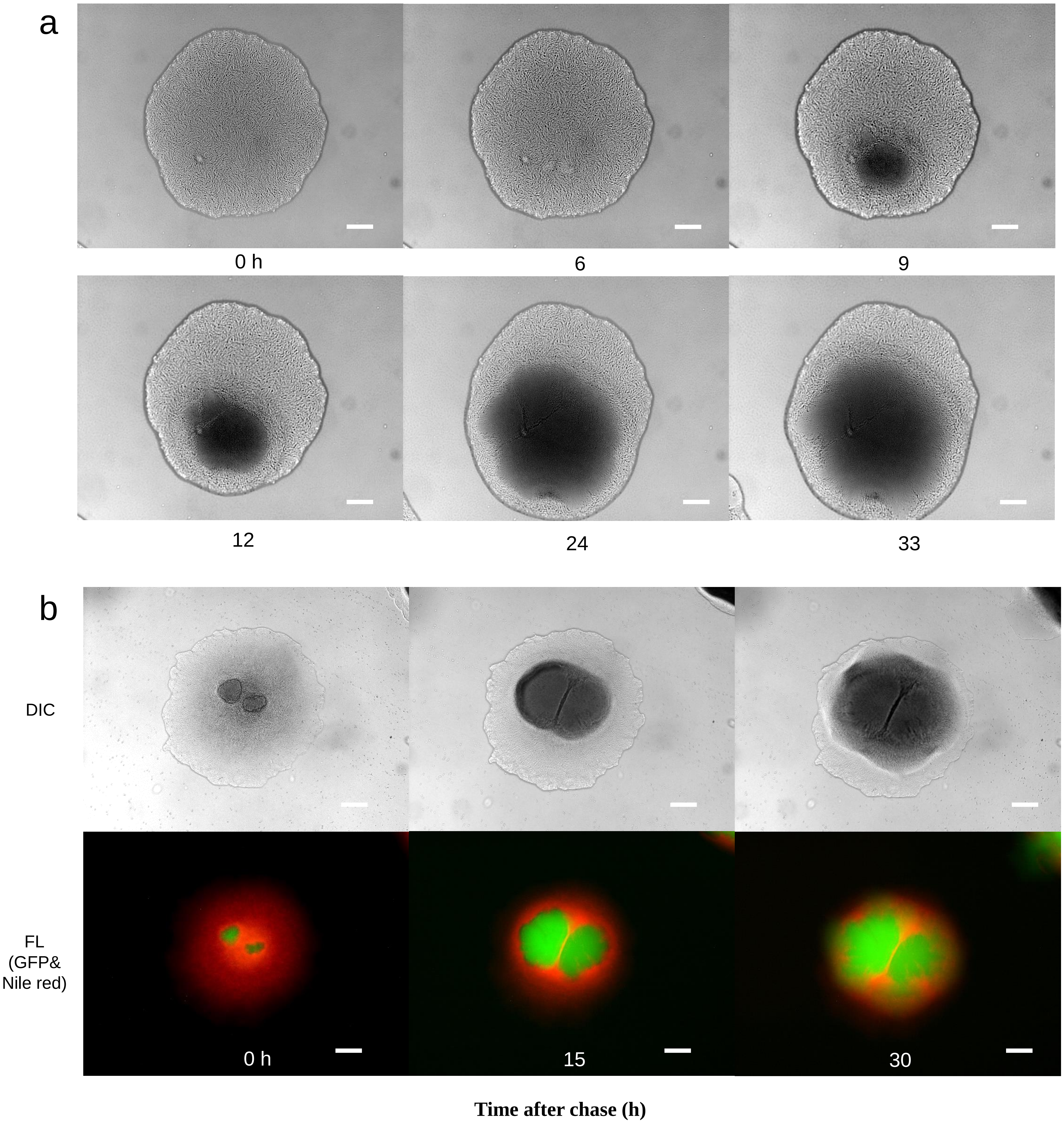

#### Supplementary Figure 9 | Growth in cargo and spawning tissue.

In a 5-day-old mature colony, coccobacillus cells start to form internally. Colony morphology varies depending on the colony density on a plate. The colony shown in the figure was cultured at a density of 200–300 CFU/plate.

**a,** Time-course images of the colony in **Supplementary Movie 4**.

**b,** Internal growth of coccobacillus cells in filamentous cargo. Nile-red dye only stained the filament cells, whereas the disorderly proliferating coccobacillus cells exhibited GFP fluorescence and turned the DIC field dark. Time after chase is shown. Bars = 100 μm.

a

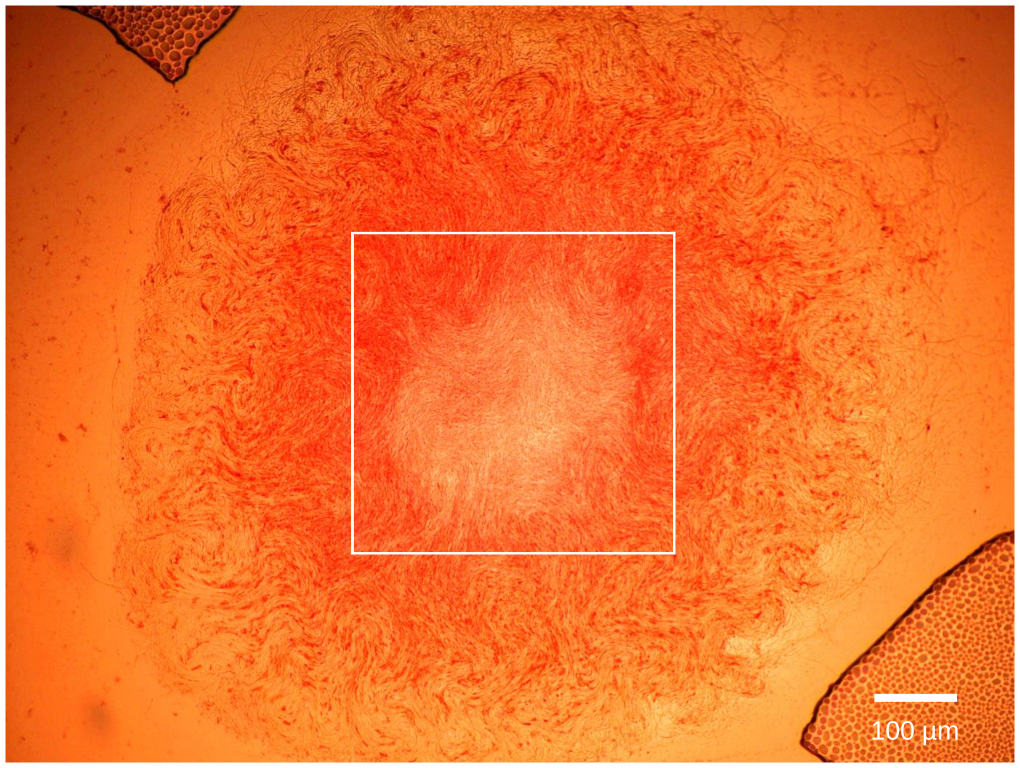

b

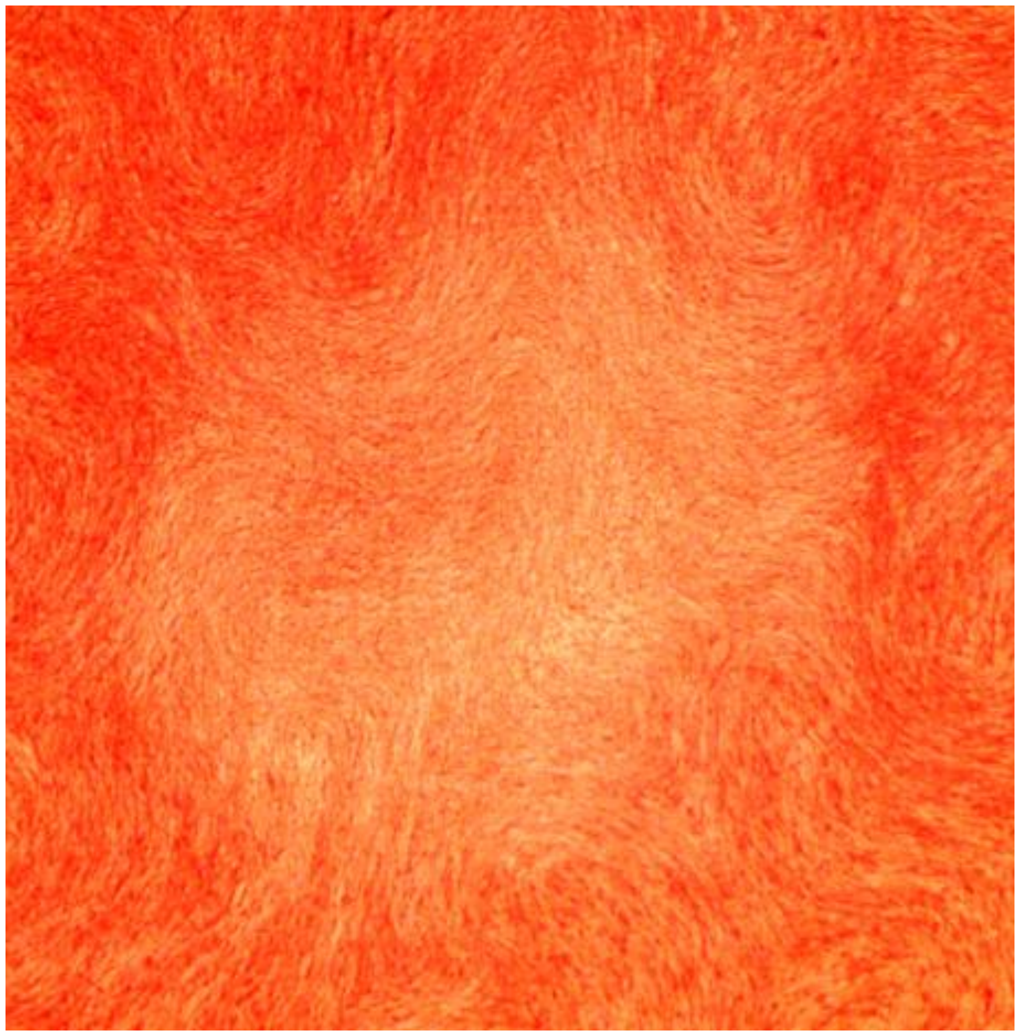

**Supplementary Figure 10 | The center area of a colony after contact with water.**

The colony was stained by fuchsine after washing.

**a**, DIC image of a fuchsine-stained colony.

**b**, A magnified image of the area indicated by the box in **a**.
The bottom cell layer sustained tightly aligned filamentous cells even after the release of coccobacillus cells.

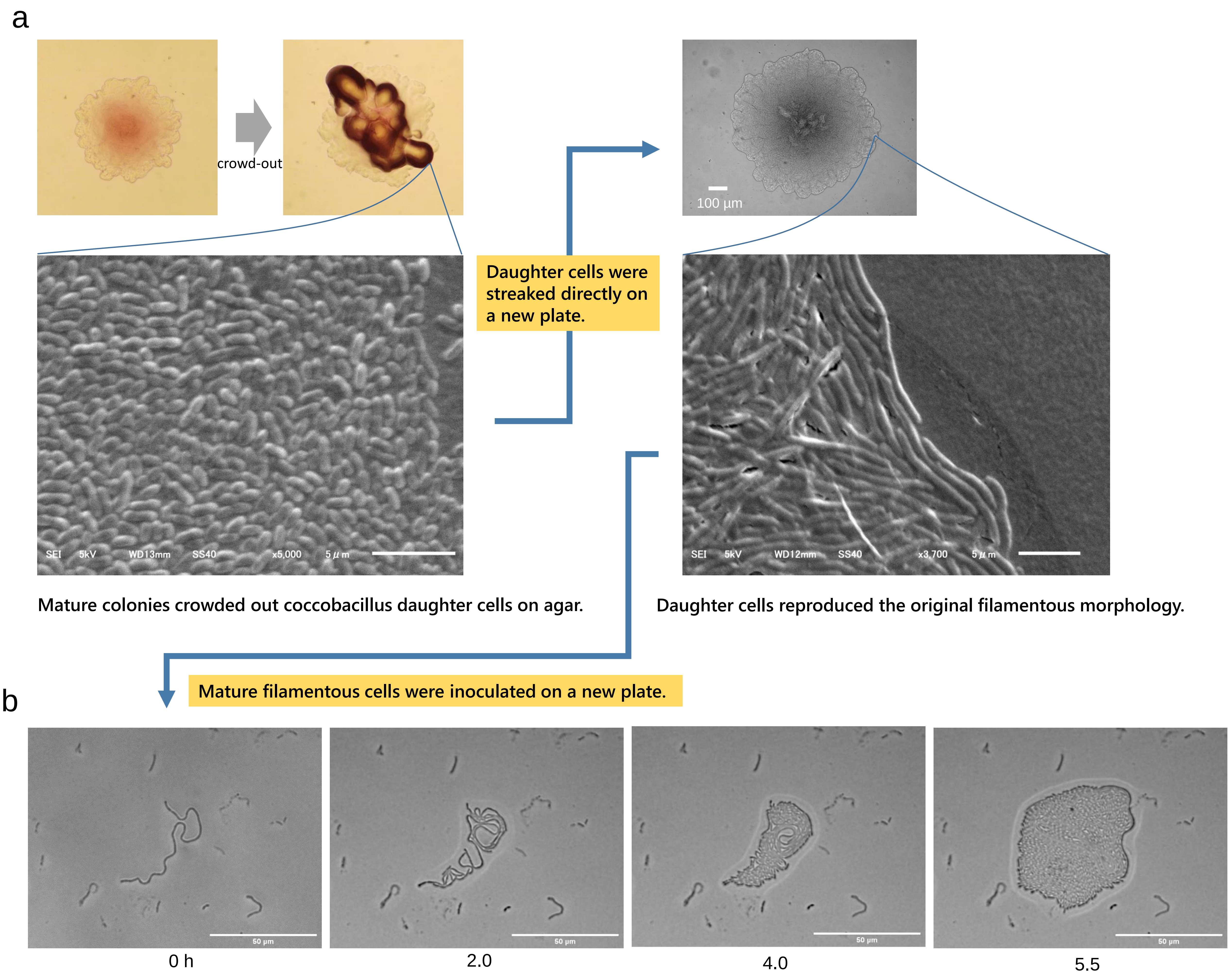

**Supplementary Figure 11 | Reversibility between filament and coccobacillus cells.**

**a**, HS-3 crowds out coccobacillus cells from mature colonies. Streaking of the daughter coccobacillus cells on a new plate showed that the coccobacilli could reproduce the original colony morphology with filamentous cells. Phase contrast images (top left) and a DIC image (top right), and SEM images (second row).

**b**, The filamentous cells were selected from a single 2-day-old colony, suspended in PBS, and inoculated on a new agar plate. The mature filament cells that survived physical damage during inoculation grew by dividing on agar. The microscopic observation was performed without a cover slip. The time after chase is shown. Bars = 50 µm. The original movie is available as **Supplementary Movie 6**.
